# Fascin in Dendritic Protrusions is Required for Synaptic Plasticity

**DOI:** 10.1101/2025.10.03.680376

**Authors:** James Q Zheng, Shuristeen Joubert, Carlos Gonzalez-Islas, Yuki Ogawa, Abhishek Poddar, Arjolyn B Penas, Enyu Liu, Peter Wenner, Kenneth R Myers

## Abstract

The Fascin family of actin-bundling proteins is best known for organizing actin filaments (F-actin) into tightly packed bundles that drive dynamic membrane protrusions such as filopodia. In neurons, fascin has been thought to primarily function in axons, as previous studies reported its absence from dendritic filopodia and spines. Here, we demonstrate that fascin is both present and functionally important in dendritic compartments. Using optimized immunocytochemistry and CRISPR-based endogenous tagging of fascin1 in cultured hippocampal neurons, we show that fascin is concentrated in developing dendritic filopodia and enriched in mature dendritic spines. Super-resolution imaging further reveals that fascin is organized into discrete nanoscale foci within spine heads, but not the spine neck. Finally, we show that CRISPR-mediated knockout of fascin1 in mature hippocampal neurons impairs synaptic potentiation, without affecting baseline excitatory synaptic transmission. Together, our findings uncover a previously overlooked aspect of actin organization in dendritic spines and establish fascin as a critical regulator of postsynaptic plasticity.

**Summary Statement:** The actin bundling protein fascin localizes to dendritic filopodia and spines, where it regulates activity-dependent synaptic plasticity.

## Introduction

Chemical synapses are specialized sites of connection that enable neuronal communication underlying sensorimotor behaviors, learning, memory, and cognition. Synapses are composed of pre- and post-synaptic structures specialized for neurotransmitter release and reception, respectively. These structures are highly plastic, undergoing both functional and structural modifications in response to activity (1–3). This includes alterations in postsynaptic surface neurotransmitter receptors and presynaptic neurotransmitter release probability (4–8). Synaptic plasticity represents the cellular basis for learning and memory (9, 10), and its disruption contributes to a wide range of neurological disorders (11–14).

Most excitatory synapses in the vertebrate brain reside on dendritic spines, which are tiny actin-rich protrusions that concentrate postsynaptic components for effective excitatory synaptic transmission (15–17). Spines undergo structural changes in shape and size during development and synaptic plasticity that are strongly associated with learning, aging, and neurodegenerative diseases (11–14). Dendritic filopodia, which act as spine and synaptic precursors, are thin actin-rich structures that sample the neuropil for axonal partners before stabilizing upon contact (18, 19). Both dendritic filopodia and spines depend upon the actin cytoskeleton for their formation, remodeling, and maintenance (16, 20–22). Unlike conventional filopodia, which contain tightly bundled unipolar arrays of parallel actin filaments, dendritic filopodia and thin spine necks contain branched and straight actin filaments of mixed polarity that do not form tight bundles (23, 24). These networks are organized and remodeled by a large number of actin binding proteins (ABPs) (25–28). During structural long-term potentiation, distinct sets of ABPs sequentially reorganize, stabilize, and consolidate the actin network in individual spines (29). However, whether additional ABPs are present in dendritic spines and contribute to the actin remodeling underlying postsynaptic plasticity remains largely unexplored.

Fascin is a highly conserved actin-bundling protein best known for generating tightly packed filamentous actin (F-actin) bundles in dynamic membrane protrusions such as filopodia and invadopodia (30–34). In neurons, fascin has been mostly studied in developing axons, where it localizes to growth cone filopodia and contributes to axon elongation and guidance (35–37). However, fascin was reported to be absent from dendritic filopodia and spines (24, 38), leading to the prevailing view that fascin plays no role in postsynaptic actin organization or plasticity. In this study, we challenge this model by showing that fascin is present and functionally important in dendritic compartments. Using a combination of confocal microscopy, super-resolution imaging, and CRISPR/Cas9 gene editing, we show that fascin is enriched in dendritic filopodia and mature spines, where it forms discrete nanoscale foci within spine heads.

Functionally, we demonstrate that CRISPR-based knockout of fascin1 in mature hippocampal neurons impairs activity-dependent synaptic potentiation without affecting baseline neurotransmission. These findings reveal an unexpected role for fascin in the modulation of the dendritic actin cytoskeleton and establish fascin as a important regulator of synaptic plasticity.

## Results and discussion

### A unique requirement for fixation methods to preserve the spatial localization of fascin

Fascin is the primary actin-bundling protein in filopodia and other cellular protrusions, yet its reported cellular distribution by immunocytochemistry has been inconsistent depending on fixation conditions. Because the localization of an actin-binding protein fundamentally shapes the interpretation of its function, resolving these discrepancies is critical for understanding fascin function in cells. This is particularly important in neurons, where the actin cytoskeleton forms highly specialized structures that support diverse neuronal functions. Previous work noted that fascin association with F-actin bundles can be detected by cold methanol fixation but not aldehyde-based fixation; however, the mechanistic basis for this fixation-dependent difference has not been elucidated (39–41). To systematically evaluate how fixation conditions affect the spatial localization of fascin in cells, we first examined mouse neuroblastoma CAD cells, which form prominent lamellipodia and filopodia when seeded on laminin substrates (42, 43). Consistent with previous findings (34, 37), cold 100% methanol (−20°C) preserved fascin enrichment in F-actin-rich lamellipodia and filopodia, as evidenced by strong fascin signals associated with F-actin bundles using multiple independent anti-fascin antibodies (Figure 1A and Figure S1). However, cold methanol fixation disrupts the binding of phalloidin to F-actin (Figure 1A), as reported previously (44, 45).

**Figure 1.**
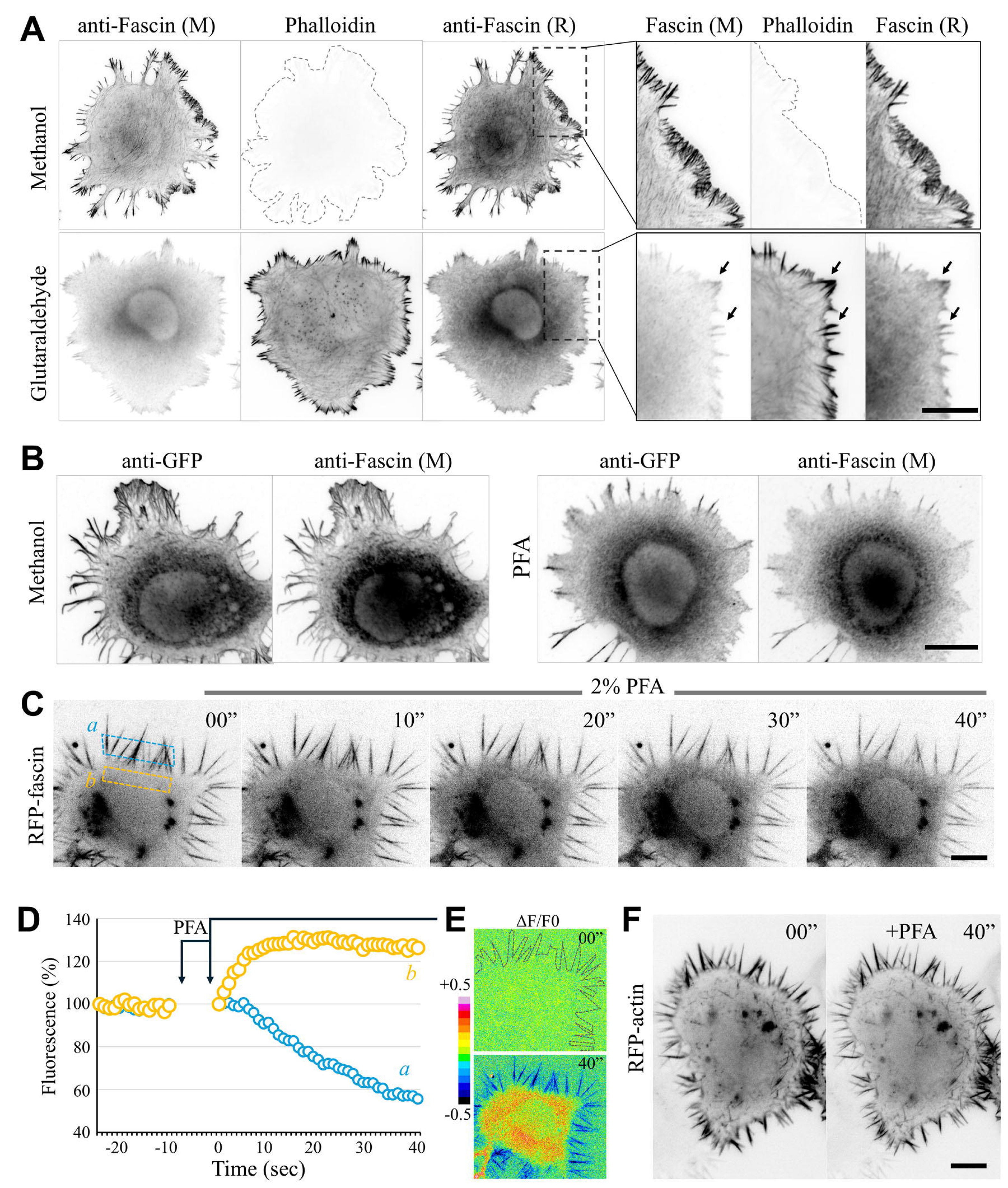
Methanol fixation is required to capture the cellular localization of fascin. (**A**) Representative immunofluorescent images of CAD cells fixed with 100% cold methanol (top row) or 2% glutaraldehyde (bottom row). Cells were stained with mouse (M) anti-fascin, rabbit (R) anti-fascin, and phalloidin-AlexaFluor546. Regions enclosed by dashed rectangles are shown at a higher magnification on the right, highlighting the actin-rich lamellipodia and filopodia. Arrows highlight weak fascin signals remaining after aldehyde fixation. (**B**) Representative images of CAD cells expressing GFP-fascin1 for approximately 24 hours and fixed with either 100% cold methanol (left panels) or 4% PFA (right panels). Cells were stained with mouse (M) anti- fascin and rabbit anti-GFP. (**C-E**) Fast time-lapse fluorescent imaging (1 frame/sec) showing live CAD cells expressing mCherry-fascin1 (referred to as RFP-fascin1) before, during, and after the perfusion of 2% PFA into the imaging chamber. The perfusion took <10 sec to complete. Times shown are in seconds with the time at 00’ indicating the end of PFA perfusion. (**C**) Representative images of a CAD cell expressing RFP-fascin1 at various time points after the PFA perfusion. The fluorescent intensities of two regions enclosed by rectangular boxes were background subtracted and quantified at every frame before and after the PFA perfusion (**D**). Due to the subtle movement of the sample during perfusion (<10 sec), fluorescence levels were not quantified until the medium was completely replaced by 2% PFA solution. The measured RFP-fascin1 fluorescence was normalized to the first frame in the plot. (**E**) Representative pseudocolor images depicting ΔF/F_0_ at 00” and 40” after PFA. The color bar indicates the colors corresponding to ΔF/F_0_ values. (**F**) Representative images of a CAD cell expressing mCherry-γ-actin (RFP-actin) at 00” and 40” after PFA perfusion. Here, the fluorescent time-lapse imaging was done at 1 frame/sec before, during, and after PFA perfusion. Given no changes in the distribution of fluorescent mCherry-γ-actin, only two time points were presented. Scale bars = 10 µm.

Conversely, fixation with either 2% glutaraldehyde or 4% paraformaldehyde (PFA) resulted in robust phalloidin staining but substantially reduced fascin signals associated with F-actin structures, rendering fascin mostly cytosolic (Figure 1A and Figure S2A).

Similarly, fixation with 50% cold methanol (24) resulted in a substantial loss of fascin signals associated with F-actin in membrane protrusions (Figure S2B). In both aldehyde- and 50% methanol-fixed CAD cells, only weak fascin signals were observed in association with thick F-actin bundles in membrane protrusions (arrows in Figure 1A and Figure S2). Similar to CAD cells, aldehyde-based fixation also diminished fascin signals associated with F-actin in nerve growth cones in culture (Figure S3). These results demonstrate that cold 100% methanol, not aldehyde-based cross-linking, is required to preserve and reveal the spatial localization of fascin to distinct F-actin structures. The substantial loss of fascin signals associated with F-actin upon aldehyde or 50% methanol fixation could lead to an underestimation of fascin enrichment in specific cellular compartments such as dendritic spines, producing incorrect interpretations about the molecular composition of these actin-based structures.

Aldehyde-based cross-linking can impair antibody detection by masking specific epitopes (46–48). To test whether this mechanism accounts for the loss of fascin signal, we expressed exogenous GFP-fascin and stained cells with both anti-GFP and anti-fascin antibodies. Aldehyde-based fixations, but not 100% methanol fixation, essentially eliminated the colocalization of GFP-fascin with F-actin bundles (Figure 1B). Critically, this loss of colocalization was observed with both antibodies, including the anti-GFP antibody known to reliably detect GFP in PFA-fixed cells (49, 50), arguing against a simple epitope-masking explanation.

To understand the potential mechanism underlying the failure of aldehyde-based fixation to preserve fascin–actin association, we performed live-cell imaging of fluorescent protein-tagged fascin1 and γ-actin during PFA fixation. To ensure that changes in fluorescence distribution reflected the behavior of the tagged proteins rather than potential differences in fluorophore-dependent responses to aldehyde fixation, we performed single-channel live imaging of RFP-tagged fascin1 and γ-actin. Imaging at 1 frame/s revealed that RFP-fascin1 (mCherry-fascin1) rapidly dissociated from the actin-rich filopodia at the cell periphery and became elevated in the cytosol immediately following perfusion with 2% PFA (Figure 1C). Quantitative measurements showed that cytosolic RFP-fascin1 fluorescence rapidly increased within the first 10 s after PFA application and reached a plateau by approximately 20 s (Figure 1D, yellow symbols). Conversely, RFP-fascin1 fluorescence in filopodia began to decline within the first 10 s and continued to decrease for approximately 40 s (Figure 1D, blue symbols). This redistribution was further illustrated by pseudocolor images depicting fluorescence changes over time (ΔF/F₀), where F_0_ is the image at time zero. After 40 sec of PFA treatment, the cell exhibited negative ΔF/F₀ values in peripheral filopodia and positive ΔF/F₀ values in the cytosol, confirming the loss and gain of RFP-fascin1 in filopodia and cytosol, respectively (Figure 1E). In contrast, RFP-γ-actin continued to label F-actin structures, with little detectable redistribution following 2% PFA fixation (Figure 1F).

Fascin exhibits rapid and dynamic binding behavior when bundling actin filaments, with a kinetic off-rate of ∼0.12 s⁻¹ (51, 52). In contrast, formaldehyde fixation proceeds through a relatively slow two-step process: an initial, rapid Schiff base formation (seconds) followed by a much slower cross-linking step requiring minutes or longer to form stable bonds (53). This temporal mismatch means that formaldehyde cannot cross-link fascin to actin filaments before fascin dissociates. Instead, formaldehyde likely inactivates fascin molecules while they are transiently unbound, trapping them in a cytosolic state, a model supported by the live-imaging data (Figure 1C-F). Together, these findings establish that rapid fixation using cold methanol (or cold acetone; data not shown) is essential for accurately visualizing fascin localization in cells, and they provide a mechanistic explanation for long-standing discrepancies in fascin localization across studies. More broadly, these results carry important implications for interpreting published data: studies that relied on aldehyde fixation or reduced methanol concentrations, to conclude that fascin is absent from specific actin-based structures, such as dendritic spines, should be re-evaluated, as the previously reported absence of fascin may reflect a fixation artifact rather than a genuine biological observation.

### Fascin is enriched in dendritic protrusions

During synaptic development, dendritic processes elaborate actin-rich dendritic filopodia that are considered precursors of postsynaptic dendritic spines (18, 19). A previous study using 50% methanol fixation concluded that fascin is absent from dendritic filopodia, unlike conventional filopodia and axonal growth cone filopodia (24). Given that 50% methanol fixation fails to preserve fascin association with F-actin (Figure S2B), we re-examined fascin localization in dendritic compartments using cold 100% methanol fixation. Since fascin is highly enriched in axonal compartments (37), we employed very low-density hippocampal cultures (10,000 cells/35mm dish) to examine dendritic fascin distribution with minimal interference from axonal signals. At DIV11, when filopodia are the predominant dendritic protrusion type, we observed robust enrichment of fascin in dendritic filopodia, colocalizing with F-actin (Figure S4A, arrowheads). We also observed fascin enrichment in dendritic protrusions at DIV13 (Figure S4B), a stage characterized by a mix of dendritic filopodia (Figure S4B, arrowheads) and immature dendritic spines that contact axons and exhibit slightly enlarged heads (Figure S4B, arrows). Spine morphology became much more apparent by DIV18, as protrusions contacting axons exhibited clearly enlarged heads enriched with fascin (Figure S4C, arrows). Fascin signals in dendritic protrusions are relatively weaker than in axonal growth cones (GC, Figure S4B), but their presence in both filopodia and immature spines is clear. Here, axons were identified by several morphological criteria, including their substantially thinner and longer appearance compared to dendrites, their relatively uniform caliber, the presence of numerous varicosities, and comparatively weak F-actin signals.

By DIV25, dendritic spines have become predominant in low-density hippocampal cultures, where fascin can clearly be detected and appears enriched in spines (Figure 2A). To confirm this observation, we examined fascin localization in mature dendritic spines in high-density hippocampal cultures (250,000 cells/35mm dish). We took advantage of sparse CRISPR/Cas9-based GFP tagging of endogenous β-actin (GFP-endo-β-actin) in high-density cultures. GFP signals were used to create a mask that removed surrounding signals from neighboring non-transduced cells, enabling clearer examination of fascin enrichment in individual dendritic spines (Supplemental figure S5). Consistent with our observations in low-density cultures, fascin is enriched in mature dendritic spines in high-density hippocampal cultures (Figure 2B). Quantitative analysis of the spine-to-shaft ratio confirmed elevated levels of fascin in spines relative to the neighboring dendritic shaft in both low-density and high-density cultures (Figure 2C). Together, our data demonstrate that fascin is not only present but also enriched in dendritic filopodia and spines, which directly contradicts the previously reported conclusion that fascin is absent from dendritic protrusions (24) and suggests a potential role for fascin in postsynaptic structure and function.

**Figure 2.**
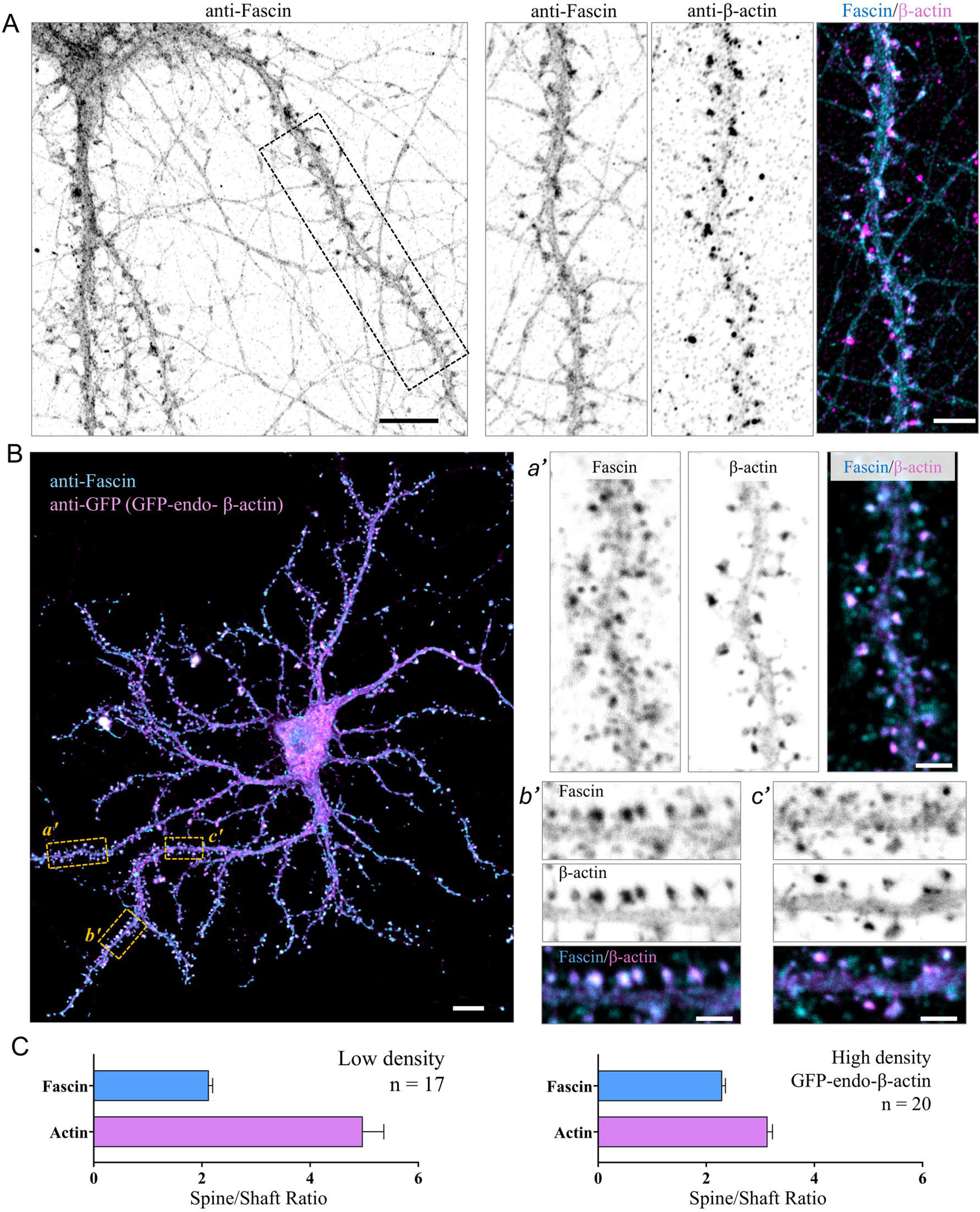
Fascin is enriched in mature dendritic spines. (**A**) Representative images of low-density cultured hippocampal neurons at DIV28 fixed with 100% cold methanol and labeled for endogenous fascin (cyan) and actin (magenta). Left scale bar = 10 µm, right scale bar = 5 µm. (**B**) Representative image of a neuron from high-density hippocampal culture expressing GFP-tagged endogenous β-actin, fixed with 100% cold methanol at DIV28, and labeled for fascin (cyan) and GFP (magenta). GFP-β-actin signal was used as mask to isolate individual neuron from neighboring cells. Right, high magnification panels of boxed regions from image on left. Left scale bar = 10 µm, right scale bars = 2 µm. (**C**) Quantification of fascin enrichment in dendritic spines. Graphs show the ratio of fascin or β-actin fluorescence intensity in spine heads compared to the intensity of an equal-sized area of the neighboring dendritic shaft. *n* = fields of view, representing >200 spines from low-density cultures and >500 spines from high density cultures, from 3 independent cultures. Error bars are ± SEM.

### Fascin in spines forms nanoscale foci

Having established that fascin is enriched in dendritic spines, we next sought to investigate whether fascin exhibits a nanoscale organization in spines. Here, we utilized CRISPR-based gene editing to fluorescently tag endogenous fascin1 at its N-terminus with smFP-V5 (54) followed by immunofluorescent labeling using anti-V5 antibodies.

Consistent with our immunostaining data, smFP-V5-endo-fascin1 signals are enriched in dendritic spines, as well as in the growth cone with filopodia (Figure 3A). Tagged endo-fascin1 signals appear to be mostly enriched in the spine heads and, interestingly, exhibit a non-uniform but punctate distribution in the spine head (Figure 3A, yellow arrows), suggesting a potentially sub-spine organization of fascin. To investigate the nanoscale distribution of untagged endogenous fascin in spines, we next performed super-resolution STED imaging. Here, high-density hippocampal cultures were subjected to GFP-tagging of endogenous β-actin via AAVs, fixed with cold 100% methanol at DIV25, and triple-labeled for GFP (for GFP-endo-β-actin), bassoon (for presynaptic active zones), and fascin. While fascin signals are clearly enriched in actin-rich dendritic spines under confocal microscopy, STED imaging reveals that fascin is distributed as discrete foci throughout the actin-rich spine head, but rarely in the spine neck (Figure 3B-C, Figure S6A). We find that individual fascin foci in spines ranged in size from 40 to 160 nm, with a mean of 97 nm (Figure 3D). This punctate pattern of fascin signals in spines contrasts sharply with the continuous fascin distribution along F-actin bundles in lamellipodia and axonal growth cones (Figure S5B-C), indicating that these nanoscale foci are not a result of fixation or immunolabeling artifacts. Rather, this organization is consistent with the absence of tight F-actin bundles in spines, suggesting that fascin may serve a distinct, bundle-independent function in regulating the dendritic spine F-actin network.

**Figure 3.**
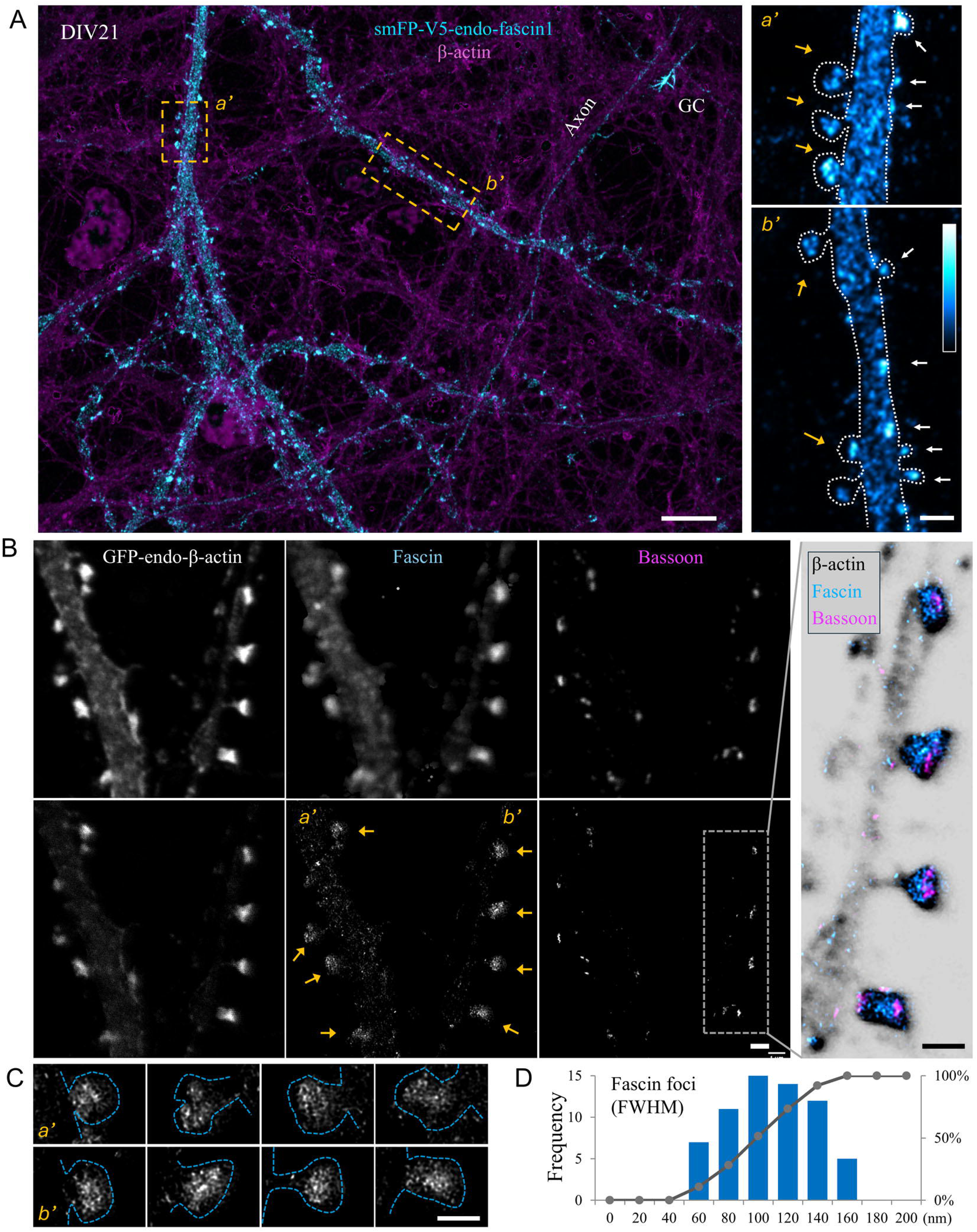
Fascin is enriched in dendritic spine heads as discrete nanoscale foci. (**A**) Representative images of hippocampal neurons at DIV21 with endogenous fascin1 tagged by CRISPR-mediated knock-in of smFP-V5 (cyan). The left panel shows a portion of the dendritic arbor of a neuron expressing smFP-V5-fascin1 (cyan) within the high-density hippocampal culture immunofluorescently labeled for actin (magenta). Fascin signals are present in dendritic arbors and enriched in dendritic spines, as well as an axonal growth cone (GC). The regions enclosed by dashed rectangles are shown in a high magnification on the right, in which fascin signals are represented using a cyan-hot look-up table. Yellow arrows indicate spines with punctate fascin distribution within the spine heads, whereas white arrows indicate small spines with fascin in their heads. Left scale bar = 10 µm, right scale bars = 2 µm. (**B**) Representative confocal (top row) and STED (bottom row) images of high-density hippocampal cultures expressing GFP-tagged endogenous β-actin, fixed at DIV25 with 100% cold methanol, and triple stained for fascin (gray), β-actin (cyan), and bassoon (magenta). GFP-β-actin signal was used as mask to isolate individual neurons from neighboring cells. Fascin appears as discrete foci within actin-rich spine heads (arrows), but not in spine necks. A portion of the dendrite (dashed rectangle) is shown on the right in a higher magnification. Here, the GFP-endo-β-actin channel is shown as inverted grayscale, overlaid with STED fascin (Cyan-Hot) and bassoon (magenta) STED signals. Scale bars = 1 µm. (**C**) High magnification images of individual spines in (B) showing the nanoscale foci of fascin (spines outlined in blue dashed lines). Scale bar = 1 µm. For the color panel in (B) and panels in (C), STED images were subjected to Abberrior’s deconvolution filter (Iterations=5, Sharpness=40, Background size=20, Background weight = 0.5). (**D**) Histogram showing the distribution of individual fascin foci widths in spine heads, grouped into 20-nm bins. Gray bars indicate the frequency of foci within each width bin. Foci width was measured as the full width at half maximum (FWHM). A total of 6 STED images comprising more than 30 spines were analyzed.

### Fascin is essential for synaptic plasticity

To test the function of fascin in dendritic spines, we utilized an AAV-based CRISPR/Cas9-mediated knockout approach to deplete fascin1 in cultured hippocampal neurons. Neurons were infected at DIV4 with AAVs expressing spCas9 and either non-targeting control gRNAs (3xgRNA-KO template) or three gRNAs targeting the coding region of FSCN1 (3xgRNA-KO-Fascin1). Double immunofluorescent labeling of methanol-fixed neurons for HA (smFP-HA) and fascin showed that most HA-positive neurons (indicating gRNA expression) exhibited essentially no fascin signal (Figure S7A). Quantification confirmed that fascin fluorescence in most HA-positive neurons was reduced to background levels, with only a small subset retaining low but detectable signal (Figure S7B), likely representing unedited or monoallelically edited cells. At the population level, however, the cultures used for western blotting also contained non-transduced neurons, which contributed to the residual fascin1 signal in the KO group (Figure S7C–D). Thus, although most individually-edited neurons showed complete loss of fascin, there remains residual signals in bulk western blot measurements which likely reflect incomplete cell transduction, rather than incomplete editing. We therefore retain the term “knockout” (KO) for this approach, while noting that at the population level it produces a substantial knockdown of fascin1. Notably, western blotting revealed a relatively late onset of fascin loss (∼12 days post-infection), consistent with a recent report of continued editing in post-mitotic cells for at least 16 days after viral sgRNA and Cas9 delivery (55). We also found that fascin expression increases gradually over the first three weeks in culture (Figure S7E), consistent with previous reports (56, 57).

We found no differences in either dendritic spine density or morphology between the control (3xgRNA-KO template) and KO (3xgRNA-KO-Fascin1) neurons (Figure 4A- D). Likewise, whole-cell patch-clamp recordings revealed no difference in the baseline frequency or amplitude of miniature excitatory postsynaptic currents (mEPSCs) between the control and KO neurons (Figure 5). This suggests that fascin1 is not required for the development of mature dendritic spines and synapses under basal conditions. To test whether fascin is required for synaptic plasticity, we used a well-established chemical LTP (cLTP) protocol in which tetraethylammonium (TEA) is applied to block potassium channels (58–60). Consistent with previous findings, brief exposure to 25 mM TEA resulted in increased spine size (Figure 4E-F) and produced robust potentiation of mEPSC frequency and amplitude in control neurons (Figure 5B-C, 5E). In contrast, fascin KO neurons showed no increase in spine size (Figure 4E-F).

**Figure 4.**
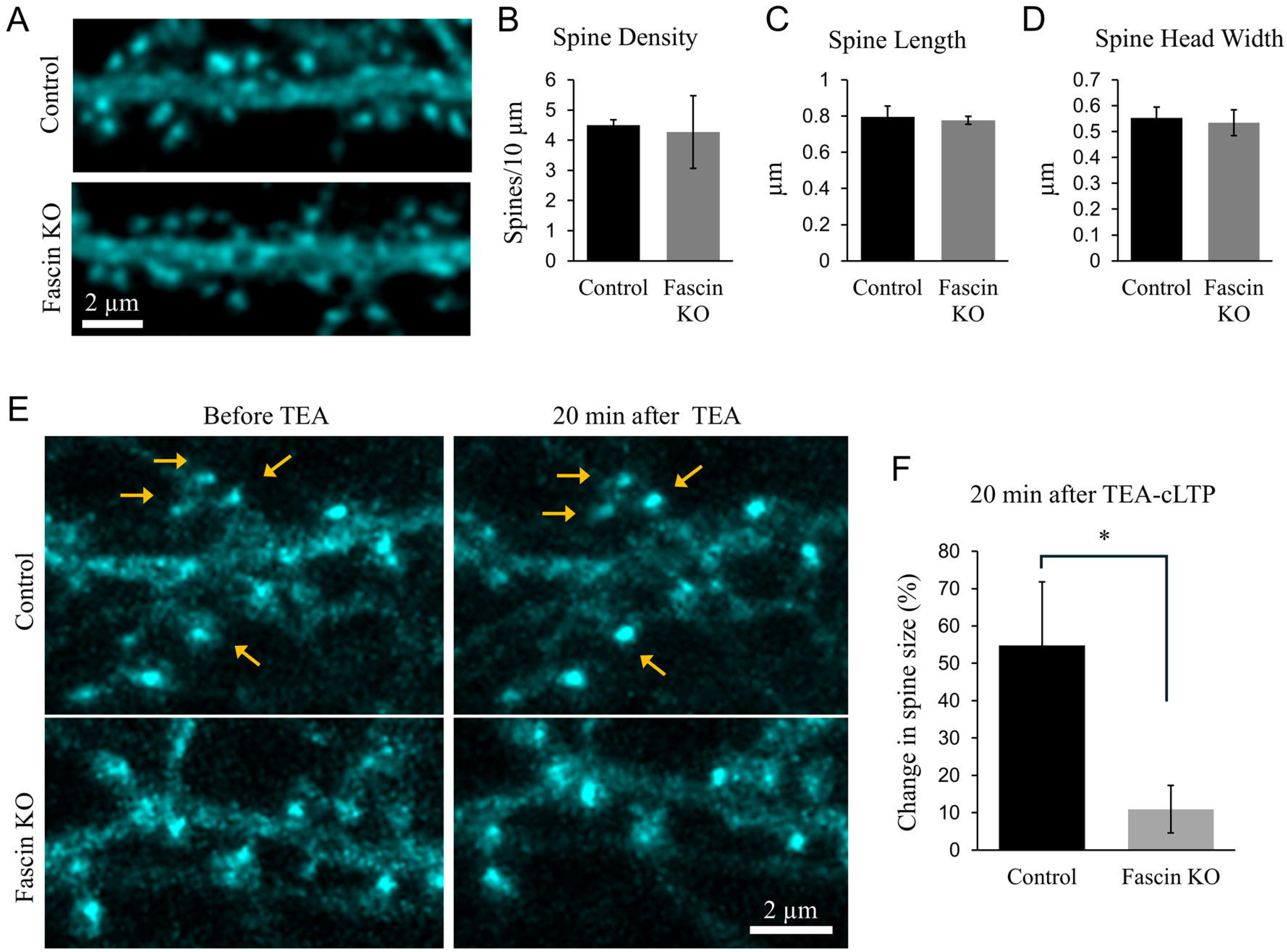
Dendritic spines at baseline and after TEA-induced chemical LTP. (**A**) Representative images of dendritic spines from control (3xgRNA-KO template) or fascin KO (3xgRNA-KO-Fascin1) high-density hippocampal neurons at DIV25-28. Co- expressed smFP is used to visualize spine morphology. Scale bar = 2 µm. (**B-D**) Plots show no differences in spine density, length, or head width between control and fascin KO cells under basal conditions (not significant by unpaired student’s *t*-test). Error bars are ± SEM. (**E**) Representative fluorescent images of dendritic regions from control and fascin KO neurons showing the spine size changes before and 20 min after TEA treatment. Arrows indicate the spines whose sizes increased at 20 min after TEA treatment. (**F**) The plot shows the changes in spine size 20 min after TEA treatment compared to that before TEA. The integrated fluorescence intensity of >150 individual spines per condition, before and after TEA treatment, from 6 independent experiments were quantified. P-values for unpaired student’s *t*-test is 0.0361. *P<0.05 (student *t*- test). Error bars are ± SEM.

**Figure 5.**
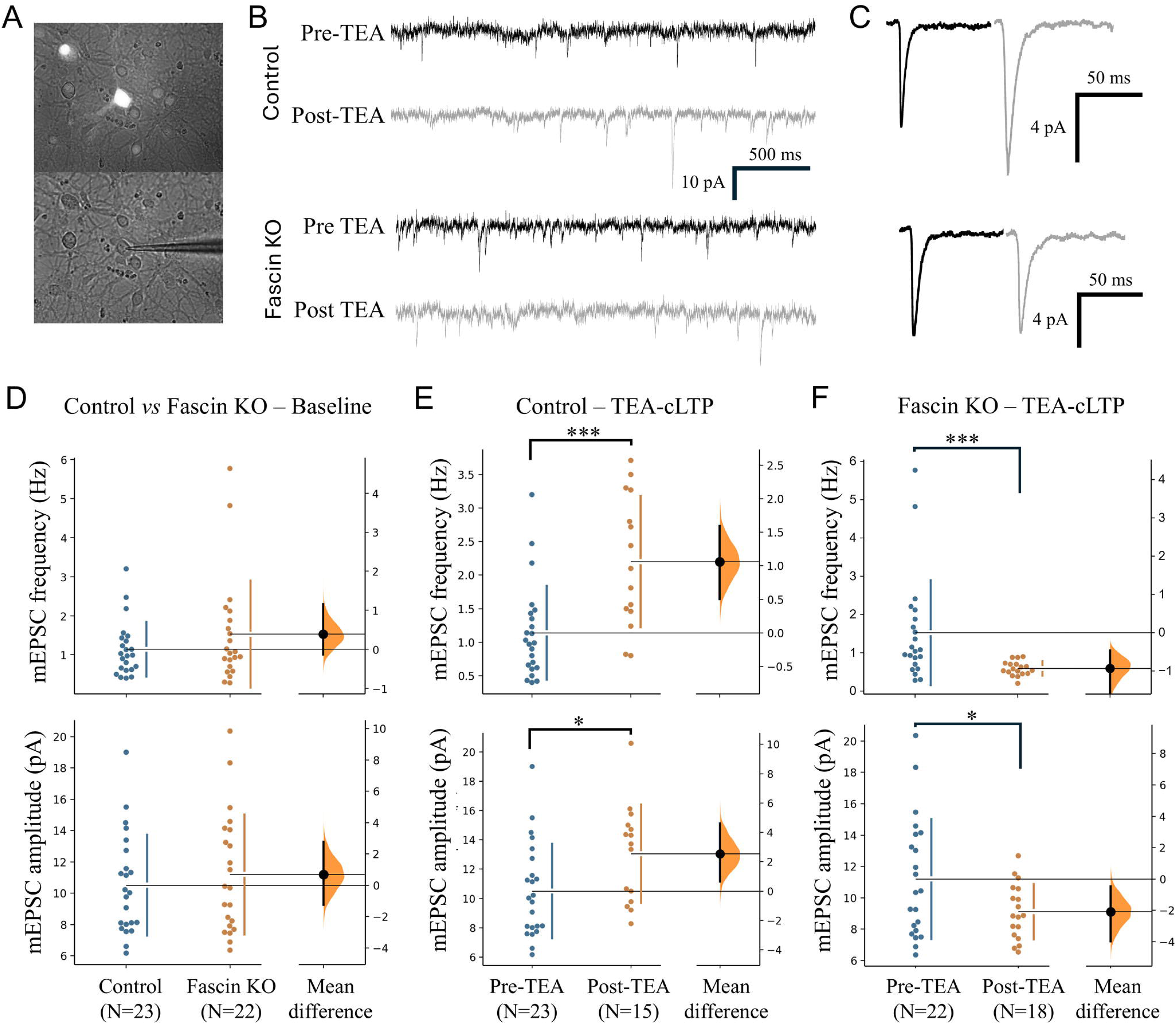
Fascin is required for synaptic potentiation in mature hippocampal neurons. (**A**) Representative images showing the identification of fluorescent neurons for patch-clamp recording. (**B**) Representative sample traces of mEPSCs from whole- cell patch-clamp recordings of control (template) and fascin KO (3gRNA) before and after application of 25 mM TEA. (**C**) Average mEPSC waveforms before (black) and after (gray) TEA application in control (top) and fascin1 KO (bottom) neurons. (**D-F**) Gardner-Altman estimation plots showing the differences in mEPSC frequency (top) and amplitude (bottom) between the control (N=23) and fascin KO (N=22) groups. (D) The baseline synaptic transmission was not affected by fascin KO, as the unpaired mean differences for the frequency and amplitude are 0.387 Hz (95% confidence interval (CI), -0.13, 1.15) and 0.694 pA (95% CI, -1.25, 2.77). P values of the two-sided permutation *t*-test on the frequency and amplitude are 0.25 and 0.52, respectively. (E) TEA-cLTP significantly increased the frequency and amplitude of synaptic transmission in the control group. The unpaired mean differences before and after TEA treatment are 1.06 Hz (95% CI, 0.503, 1.6) and 2.55 pA (95% CI, 0.639, 4.61). (F) TEA-cLTP did not increase, but decreased, the frequency and amplitude of synaptic transmission in the fascin KO group. The unpaired mean differences before and after TEA treatment are - 0.935 Hz (95% CI, -1.59, -0.466) and -2.08 pA (95% CI, -3.96, -0.478). ***: P<0.001, * P<0.05 (two-sided permutation *t*-test).

Interestingly, rather than increasing, both mEPSC frequency and amplitude decreased after TEA treatment (Figure 5B-C, 5F). These results identify fascin as a key actin regulator required for activity-dependent synaptic remodeling.

The actin cytoskeleton is highly enriched in postsynaptic spines where it not only supports the distinct spine shapes but also provides a scaffold for the spatiotemporal organization of postsynaptic receptors and signaling components (21, 61). Importantly, postsynaptic remodeling during learning and memory depends on highly regulated actin dynamics, and its disruption is strongly associated with neurological conditions (4, 16, 62). Despite tremendous progress in understanding actin-based cellular activities, we still lack a detailed picture of actin organization and dynamic regulation in synaptic compartments. In this study, we establish fascin as a key actin regulator of postsynaptic structural and functional plasticity, revising the prevailing view that fascin is neither present nor functionally relevant in dendritic compartments. These findings are consistent with recent studies identifying fascin1 as a target for spine formation and memory consolidation (63, 64).

How fascin regulates the spine F-actin network remains unknown. Our STED imaging reveals that fascin forms discrete nanoscale foci in spine heads rather than the continuous distribution seen along F-actin bundles in filopodia and lamellipodia. While dendritic spines appear to lack extensive uniformly aligned F-actin bundles, previous studies have identified distinct actin filament structures within spines, including F-actin bundles, actin-polymerization hotspots, and nanoscale actin foci (24, 65, 66) In the current study, we visualized GFP-tagged endogenous β-actin, which highlights both the filamentous and monomeric pools of actin in spines. Our previous study shows that dendritic spines contain a pool of monomeric actin that plays a role in postsynaptic plasticity (67). The presence of monomeric GFP-β-actin in spines can occlude the distinct F-actin structures that were previously visualized by STED imaging using a F-actin-specific probe, such as LifeAct (65). Determining which F-actin structures fascin engages within these foci would be an important next step. Such structural characterization is typically achieved through electron microscopy approaches, particularly platinum replica electron microscopy, as employed previously to examine actin architecture in spines (24). However, these EM approaches require strong chemical fixation and detergent extraction steps that, as we demonstrate here, displace fascin from F-actin structures. This technical limitation likely explains why fascin was not detected in dendritic protrusions in earlier ultrastructural studies. As we have shown, aldehyde or reduced-methanol fixation caused a substantial loss of fascin molecules associated with F-actin structures in the cell periphery but retained a tiny amount of fascin signals associated with the thickest, most stable F-actin bundles. Compared to typical filopodia of axonal growth cones and motile cells that contain relatively high amounts of tightly packed F-actin bundles, dendritic filopodia and spines contain a much lower amount of bundled actin and associated fascin. This difference likely explains why earlier work detected fascin on axonal growth cones but not dendritic filopodia within the same field of view. Future advances in cryo-electron microscopy of neuronal samples may ultimately enable visualization of fascin-actin interactions in spines under near-native conditions.

Given the lack of extensive tight actin bundles in spine heads, it is plausible that fascin adopts a distinct organization within the mostly branched F-actin network to function in a complementary, but non-overlapping, manner from other actin-bundling proteins in spines (52, 68). For example, α-actinin-mediated crosslinking of actin filaments is important for proper spine development and maintenance, as well as for linking actin to the postsynaptic density (69, 70). However, unlike α-actinin, which localizes throughout dendritic spines (71, 72), the appearance of fascin in discrete nanoscale foci specifically within spine heads (Figure 3B) suggests a role in fine-tuning actin dynamics within the densely branched actin network of the spine head. Recent studies indicate that fascin binding to F-actin strongly impairs cofilin-mediated actin turnover, even without forming thick actin filament bundles (73), raising the possibility that fascin could regulate actin turnover within the branched F-actin network of dendritic spines. In this model, a small number of fascin molecules engaging neighboring filaments within the branched spine network could locally modulate cofilin-mediated regulation of actin dynamics, which is important not only for control of spine structural remodeling but also for the regulation of synaptic receptor surface insertion (60). Recent evidence in non-neuronal contexts also supports a role for fascin in the regulation of branched actin networks (73, 74).

It is notable that TEA-cLTP produced a decrease, rather than simply a failure of potentiation, in both mEPSC frequency and amplitude in fascin KO neurons. One possibility is that the activity-dependent calcium influx triggered by cLTP induction drives cofilin-dependent actin remodeling in spines that, in the absence of fascin-mediated stabilization, results in a net loss of synaptic efficacy rather than potentiation. Although our conclusion on the fascin role in postsynaptic plasticity is based on TEA-induced chemical LTP, this actin-based mechanism is likely to function in other forms of synaptic plasticity, though this requires further investigation. Additionally, because changes in mEPSC frequency can reflect presynaptic alterations, and fascin is known to be present in developing axons (37), it will be important to determine whether fascin also contributes to actin organization in mature presynaptic terminals. It should be noted that a change in mEPSC frequency during plasticity can also be expressed postsynaptically through a change in the number of functional (AMPAR-containing) synapses, which can raise or lower the frequency without any change in presynaptic release. For example, the activity-dependent insertion of AMPARs at synapses that lack them (silent synapses) increases the number of functional synapses and thereby the mEPSC frequency (60, 75, 76). Future studies are needed to elucidate the precise mechanism by which fascin engages with F-actin in postsynaptic compartments, whether it also functions presynaptically, and how these roles contribute to synaptic plasticity.

### Experimental Procedures

#### Cell culture

Dissociated primary hippocampal neuron cultures were prepared as previously described (60). Briefly, 18.5-day-old embryos were collected from timed-pregnant Sprague Dawley rats (Charles River Laboratories), and hippocampi were dissected in ice-cold Hank’s Balanced Salt Solution (HBSS; 14175095; Gibco). Hippocampi were pooled together, trypsinized for 12 minutes (25200056; Gibco), briefly incubated in 20% fetal bovine serum (A5670601; Gibco), and plated on 25 mm acid-washed glass coverslips coated with 100 μg/ml poly-L-lysine (P2636; Sigma-Aldrich) at a density of approximately 300,000 cells per 35 mm dish. Hippocampal neurons were cultured in Neurobasal medium (21103049; Gibco) supplemented with B-27 (17504044; Gibco), penicillin/streptomycin (30002CI; Thermo Fisher Scientific), and GlutaMax (35050061; Gibco), and fed ½ volume of fresh medium every 7 days post-plating. Animal care and use was conducted following National Institutes of Health guidelines, and procedures were approved by the Institutional Animal Care and Use Committee at Emory University.

Low-density hippocampal cultures were prepared using a feeder layer of cortical neurons according to the previously published method (77). Here, hippocampal neurons were plated on poly-L-lysine-coated 25 mm glass coverslips at a density of 25,000 cells per well in a 6-well culture plate. In a separate 6-well culture plate, several small paraffin wax dots ∼2 mm in diameter and ∼1 mm in height were planted on the bottom of each well, serving as spacers. After coating the bottom of each well with 100 μg/ml poly-L-lysine, cortical neurons from the same embryonic brains were plated at a density of 400,000 cells per well. After 24 h in culture, the coverslips with low-density hippocampal neurons were removed from their plate and placed in the 6-well plate of cortical neurons with the hippocampal neurons facing down towards the cortical neurons. Weekly ½ volume medium changes were carried out as described above.

The mouse neuroblastoma cell line, Cath.-a-differentiated (CAD) (78), was maintained in DMEM/F12 (10-092-CV; Corning) supplemented with 8% fetal bovine serum (Atlanta Biologicals) and Penicillin/Streptomycin. To obtain CAD cells with elaborated lamellipodia and filopodia, we re-suspended the cells and plated them on laminin-coated (20 µg/ml) glass coverslips for 2-3 hours at 37°C before fixation/immunofluorescence or live cell imaging (79, 80).

#### Transient transfection and AAV-based transduction

Fluorescent protein-tagged proteins were transiently expressed in CAD cells using X-tremeGENE HP (06366244001; Roche) DNA transfection reagent (79, 80). GFP-fascin1 and mCherry-fascin1 were based on YFP-fascin1 described previously (34) and GFP- or mCherry-γ-actin constructs were described previously. (67).

To endogenously tag fascin1 in cultured hippocampal neurons, we utilized the Homology-Independent Universal Genome Engineering (HiUGE) method (81, 82) and adeno-associated virus (AAV) to tag the N-terminus of Fascin1 in cultured rat hippocampal neurons. Custom single-stranded DNA oligonucleotides encoding guide RNAs (gRNAs) targeting the N-terminus (5’-TTGGCGGTCATGGTGGCGGA) of Fascin1 were synthesized, annealed, and subcloned into the pAAV-pHiUGE-smFP-V5 donor-ORF0 plasmid (Addgene #200380) after SapI restriction enzyme digestion (83). Here, we used spaghetti monster fluorescent proteins (smFPs), which are an engineered family of GFP-like fluorescent proteins carrying numerous copies (typically 10–15) of a single small peptide epitope (e.g., V5 or HA), enabling the tagged protein to be detected both by the smFP fluorescence and by strong antibody-based signal amplification for localizing even weakly expressed proteins (54). β-actin was endogenously tagged with EGFP using pAAV-pHITI-Actb-GFP as described (83) to identify and visualize sparsely distributed edited neurons for high- and super-resolution imaging of fascin in dendritic spines.

To knockout fascin1, we employed a previously described (84) AAV-mediated multiplexed gRNA system. We inserted the following 3 gRNAs targeting Fascin1 into pAAV-3xgRNA-KO (Addgene #240310): TCTACCGCGATCTCGTCGGC, GCGACCTTCGCAGTCGCGAA, and GCACCCACACGGGCAAGTAC. As the control, we used the pAAV-3xgRNA-KO-template which encodes three non-targeting gRNAs: CTGTCTTCATCATGGCCGAC, GTTCGCATTATCCGAACCAT, TAAGCGTCGCAAGAAGACGG. Both pAAV-3xgRNA-KO and pAAV-3xgRNA-KO- template also co-express a bright smFP-HA under the control of the hSyn1 promoter for easy identification of transduced neurons. To express Cas9, we used a modified version of pAAV-pMecp2-SpCas9-spA (Addgene #60957) lacking the HA-tag. The following high-titer AAV particles were produced with the PHP.S serotype by VectorBuilder (Chicago, IL): pAAV-pHiUGE-smFP-V5-Fascin1, pAAV-pHITI-Actb-GFP, pAAV- 3xgRNA-KO template, pAAV-3xgRNA-KO-Fascin1, and pAAV-pMecp2-SpCas9. For transduction of cultured neurons, we added AAV-Mecp2-SpCas9 (MOI = 50,000) together with either pHiUGE-smFP-V5-Fascin1, HITI-Actb-GFP, 3xgRNA-KO template, or 3xgRNA-KO-Fascin1 (MOI = 50,000). We applied AAV particles to DIV4 neurons for 48 hours before washout and maintained the cells until DIV21 or later for imaging and electrophysiology.

#### Electrophysiology

Whole-cell voltage clamp recordings were obtained from DIV21-23 rat hippocampal neurons expressing bright smFP-HA using an AxoPatch 200B amplifier (Molecular Devices), controlled by pClamp 10.1 software (Molecular Devices), low pass filtered at 5 KHz on-line and digitized at 20 KHz. Tight seals (>2 GΩ) were obtained using thin-walled boro-silicate glass microelectrodes pulled to obtain resistances between 7 and 10 MΩ. The intracellular patch solution contained the following (in mM): potassium-gluconate 125, KCl 10, Tris-phosphocreatine 5, Mg-ATP 3, Na_2_-GTP 1.5, HEPES 10, MgSO_4_ 2, CaCl_2_ 0.1, EGTA 0.5. The pH was adjusted to 7.4 with KOH. Osmolarity was measured between 280-300 mOsm. The recording chamber was mounted on an Axiovert 100 Microscope (Karl Zeiss Microscopes) and continuously perfused (5 ml/min) with oxygenated Artificial Cerebral-Spinal Fluid (ACSF) recording solution containing the following (in mM): NaCl 126, KCl 3, NaH2PO4 1, CaCl_2_ 2, MgCl_2_ 1, HEPES 10 and D-glucose 25. The pH was adjusted to 7.4 with NaOH. Miniature excitatory postsynaptic currents (mEPSCs) were isolated using 1 µM TTX (14964-1; Cayman) and 5 µM Gabazine (1262; Bio-Techne) in ACSF to block action potentials and GABAergic mIPSCs, respectively. Membrane potential was held at -70 mV and recordings were performed at 27 C. For chemically induced LTP, recording solution was switched to oxygenated LTP-induction solution containing (in mM): tetraethylammonium chloride (TEA-Cl) 25, CaCl_2_ 5, MgCl_2_ 0.1, NaCl 97, KCl 3, NaH_2_PO_4_ 1, HEPES 10 and D-glucose 25 for 10 min. Series resistance during recordings varied from 15 to 20 MΩ and was not compensated. Recordings were terminated whenever significant increases in series resistance (> 20%) occurred. Analysis of mEPSCs was performed blind to condition with MiniAnalysis software (Synaptosoft) using a threshold of 5 pA for mEPSC amplitude. Estimation statistics were used. 5000 bootstrap samples were taken; the confidence interval was bias-corrected and accelerated. The p values reported are the likelihoods of observing the effect sizes if the null hypothesis of zero difference is true. For each permutation p value, 5000 reshuffles of the control and test labels were performed (85).

#### Antibodies and immunofluorescent labeling

The following antibodies were used: rabbit polyclonal anti-GFP (Invitrogen, #A11122; 1:1000), mouse monoclonal anti-α-tubulin (clone DM1A, Sigma, #MABT205; 1:1000), mouse monoclonal anti-fascin (clone 55K2, Millipore Sigma, #MAP3582; 1:1000), rabbit monoclonal anti-fascin (Abcam, #EP5902, KO validated; 1:500), sheep polyclonal anti-fascin (R&D Systems, #AF7745, KO validated; 1:1000), mouse monoclonal anti-bassoon (clone SAP7F407, Enzo, #ADI-VAM-PS003; 1:500), and mouse monoclonal anti-V5 (clone SV5-Pk1, Invitrogen, #R960-25; 1:200). Mouse anti-fascin antibody was generated using purified full-length human fascin, and recognizes fascin from mouse, rat, and human. Rabbit anti-fascin antibody was generated using amino acids 200-400 from recombinant human fascin and recognizes fascin from human and mouse (predicted to recognize rat). Sheep anti-fascin antibody was generated using full-length human fascin1 and it detects human, rat, and mouse fascin. Triple channel immunofluorescent staining of cultured CAD cells using these three anti-fascin antibodies was found to label the same F-actin structures, with quantitative analysis showing a remarkably high degree of colocalization as indicated by Pearson Correlation Coefficients, Manders’ Colocalization Coefficients, and Costes’ P-values (Supplemental figure S1).

For confocal imaging, Alexa Fluor-conjugated (AF488-, AF546-, and/or AF647) secondary antibodies raised in Donkey were diluted (1:500) in phosphate-buffered saline (PBS) containing 2% donkey serum for 1.5 hours. For STED imaging, long Stokes-shifted Abberior STAR 460L-conjugated goat anti-rabbit, STAR ORANGE-conjugated goat anti-mouse, and STAR RED-conjugated donkey anti-sheep secondary antibodies were used. Due to the cross-reactivity of goat and sheep IgGs, we employed a two-step labeling approach. Cells labeled with three primary antibodies (mouse, rabbit, and sheep) were first incubated with STAR RED-conjugated donkey anti-sheep antibodies for 1.5 hours at room temperature, followed by three washes with PBS, PBS- 0.2% Tween-20, and PBS, respectively. Cells were then incubated with both STAR 460L and STAR ORANGE-conjugated goat antibodies for 1.5 hours at room temperature, followed by three washes, and mounted in ProLong Gold antifade mount (Thermo Fisher Scientific, #P36930) for over 2 days before STED imaging.

The optimal fixation method for immunofluorescent labeling of fascin is the use of 100% cold methanol. Briefly, cultured hippocampal neurons were rapidly fixed with 100% cold anhydrous methanol (Sigma; 322415) for 10 min at -20°C, followed immediately by rehydration in PBS for 10 min, a brief rinse with PBS containing 0.1% Triton X-100, and three additional washes in PBS. Cells were blocked in PBS with 2% normal donkey serum (Sigma; D9663) for 1 hour at room temperature. Neurons were incubated in a humidified chamber overnight at 4°C with primary antibodies diluted in PBS with 2% donkey serum. Cells were washed with PBS, PBS-0.05% Tween 20, and PBS (5 min each), followed by incubation with the mixture of secondary antibodies for 1.5 h at room temperature. After three washes with PBS, PBS-Tween20, and PBS, the coverslips were mounted on glass slides using ProLong Gold antifade (Invitrogen; P36930).

#### Microscopy and imaging

Laser scanning confocal microscopy was performed on a Nikon C2 confocal unit with four laser lines (405 nm, 488 nm, 561 nm, and 610 nm), three detectors, and either 10X/NA0.5 or 60X/NA1.4 Plan Apo objectives (Nikon Instruments Inc, Melville, New York, USA). Confocal image acquisition was done using Nikon Elements software.

Stimulated Emission Depletion (STED) super-resolution imaging was performed using an Abberior Facility Line STED system (Abberior Instruments GmbH, Göttingen, Germany) housed on an Olympus IX83 inverted microscope (Evident, Tokyo, Japan) with a 60X/NA1.4 oil immersion objective (UPLXAPO, Olympus, Japan). The STED system is equipped with four excitation lasers (405 nm, 20 mW continuous-wave; 485 nm, 1 mW pulsed; 561 nm, 200 µW pulsed; and 640 nm, 1 mW pulsed) and two pulsed STED depletion lasers (595 nm, 400 mW and 775 nm, 2750 mW). Using the long-Stokes-shift fluorophore Abberior STAR 460L together with STAR ORANGE and STAR RED, a single 775-nm pulsed STED depletion laser was used for simultaneous three-channel STED imaging with 485-, 561-, and 640-nm excitation. For these experiments, the 775-nm STED laser was typically operated at 15% of its maximum power. The excitation laser powers and total pixel dwell times for STAR 460L, STAR ORANGE, and STAR RED were 20% and 25 µs, 30% and 25 µs, and 20% and 25 µs, respectively.

Image acquisition was performed using iMSPECTOR software (Abberior, Germany), and post-acquisition image processing and quantification were performed using FIJI ImageJ (NIH, Bethesda, MD). All quantification was performed on the original, unprocessed images without image enhancement. For representative images, background subtraction and linear contrast adjustment were applied uniformly across experimental conditions for display purposes only and were not applied to images used for quantification. For quantification of nanoscale fascin foci, individual foci within the actin-defined spine heads were identified as local intensity maxima above background, and their size was determined from intensity line profiles using the full width at half maximum (FWHM).

Manual spine analyses were performed using ImageJ software (NIH). The dendritic branch length was measured using the segmented line measurement tool and each spine was marked with multi-point. Using straight line tool, spine width was measured at the maximum width of the spine tip, and spine length was measured from the spine tip to the dendritic shaft (86). Spine measurements were taken from at least two dendritic branches per neuron and made by an observer blinded to experimental conditions.

#### Western blots

Rat hippocampal neurons were lysed directly in 1× Laemmli sample buffer at indicated timepoints, then boiled for 5 min, and vortexed for 5 min. Equal volumes of cell lysates were loaded on mini-Protean 12% Tris-glycine acrylamide gels (#4561043, Bio-Rad) and transferred to nitrocellulose. Membranes were blocked with 5% milk in PBS-Tween (0.2%) for 1 h, then incubated overnight at 4°C with rabbit anti-Neuronal Class III β-tubulin polyclonal (#PRB-435P, Covance; 1:2000) and mouse anti-fascin (1:5000). Membranes were washed 3x in PBS-Tween then incubated in Alexa Fluor 568– conjugated anti-rabbit (Invitrogen) and IRDye800-conjugated (LiCor) anti-mouse secondary antibodies for 1 h at room temperature. Membranes were washed 3x in PBS-Tween and then visualized using a ChemiDoc MP imaging system (Bio-Rad, Hercules, CA). Quantification was done using ImageJ’s Gel Analyzer.

## Data Availability

The data supporting the findings of this study are available from the corresponding author, James Zheng, upon reasonable request. The original, unprocessed images and the data underlying the quantitative analyses presented in the figures are available upon request.

## Acknowledgements

We would like to thank Dr. Matthew N Rasband at Baylor College of Medicine for facilitating and supporting the development of the CRISPR constructs and plasmids for this study. This research is supported in part by research grants from the National Institutes of Health to JQZ (MH133798), and a Whitehall Foundation research grant to KRM (2023-08-069). This work was also supported by the Emory University Emory Integrated Cellular Imaging Core Facility (RRID:SCR_023534). The content is solely the responsibility of the authors and does not necessarily reflect the official views of the National Institutes of Health.

## Author contributions

JQZ conceptualized and initiated the project, as well as designed and conducted most of the experiments and data analysis. SJ performed the imaging and biochemical analyses on different fascin antibodies and knockout efficacy. CGI and PW helped with the electrophysiology recordings. YO helped with the CRISPR design and validation of fascin KO and KI constructs. AP helped with spine analysis. ABP performed fascin immunostaining. EL helped with spine analysis of the control and fascin KO groups. KRM helped with the overall project planning, experimental design, molecular biology, and image analysis, and worked with JQZ on the manuscript writing.

