## Supplemental figures S1-S7 for "Fascin in Dendritic Protrusions is Required for Synaptic Plasticity"

Zheng et al. “Fascin in Dendritic Protrusions Is Required for Synaptic Plasticity”

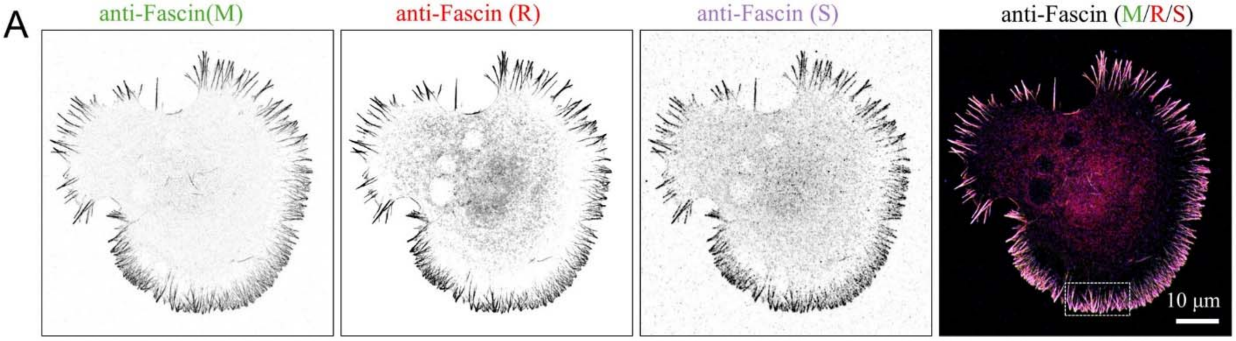


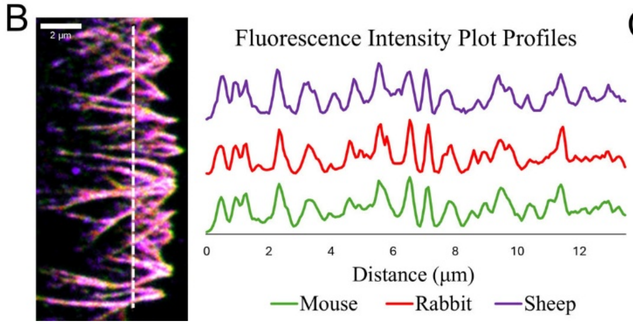


C

| Metric | Mouse and Rabbit | Mouse and Sheep | Rabbit and Sheep |
|---|---|---|---|
| Manders M1 | 0.964 | 0.997 | 0.872 |
| Manders M2 | 0.983 | 0.821 | 0.998 |
| Pearson’s R | 0.89 | 0.69 | 0.75 |
| Costes P-value | 1.00 | 1.00 | 1.00 |

**Figure S1. Fascin localization after methanol fixation is detected with multiple antibodies.** (**A**) Representative images of CAD cells fixed with 100% cold methanol (−20 °C) and stained with three independent anti-fascin antibodies (raised in mouse (M), rabbit (R), and sheep(S)). (**B**) Left, magnified image from the boxed region of (A), showing essentially the same fascin enrichment pattern in lamellipodia and filopodia using three independent antibodies. Right, fluorescence intensity profiles corresponding to the dotted white line from image on the left. (**C**) Quantification of colocalization, analyzed for each pairing of anti-fascin antibodies.

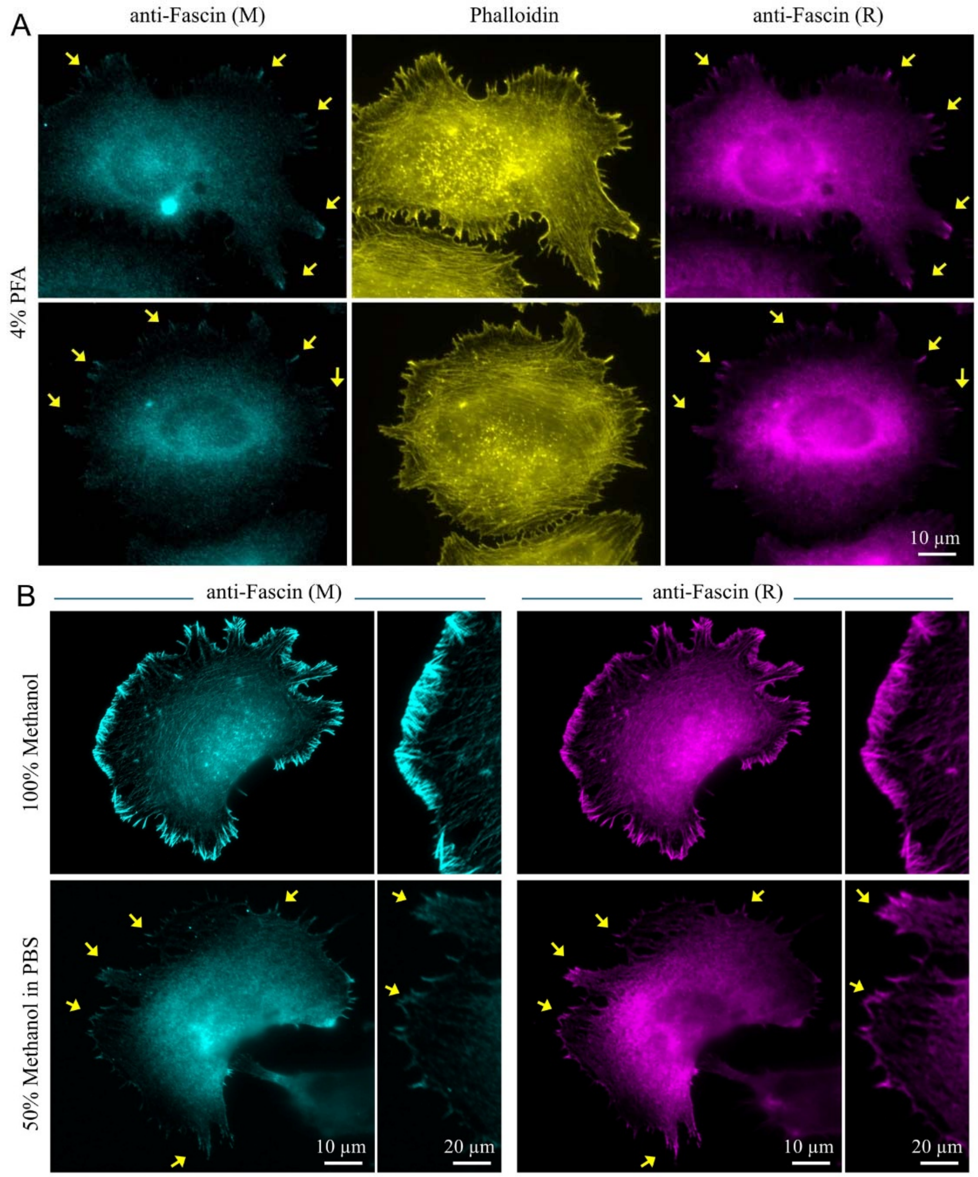


**Figure S2. Loss of F-actin associated fascin signals in CAD cells fixed with paraformaldehyde or 50% methanol.** (**A**) Representative immunofluorescent images of CAD cells fixed with 4% paraformaldehyde (PFA) stained with mouse (M) anti-fascin (cyan), fluorescent phalloidin (yellow), and rabbit (R) anti-fascin (magenta). Arrows indicate weak fascin fluorescence associated with a few thick F-actin bundles in filopodial protrusions. (**B**) Fixation with 50% methanol compromises fascin labeling, compared to fixation with 100% methanol. Representative images of CAD cells fixed with either 100% or 50% cold methanol (−20 °C) and stained with mouse (M) anti-fascin and rabbit (R) anti-fascin antibodies. Arrows indicate weak fascin immunofluorescent signals associated with a few thick F-actin protrusions in 50% methanol-fixed cells.

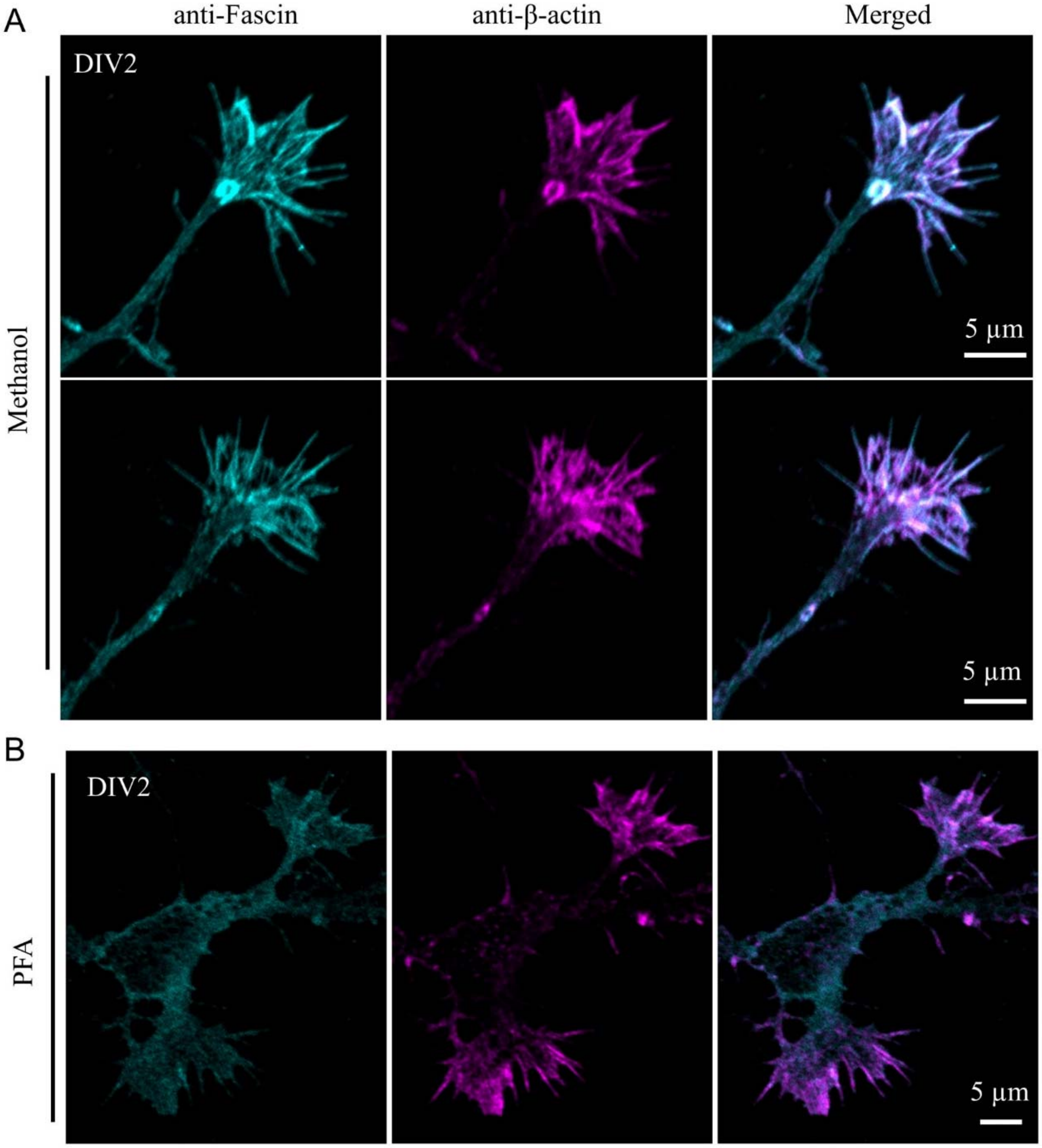


**Figure S3. Fascin signals in nerve growth cones with different fixation methods.** (**A**) Representative images of nerve growth cones from DIV2 hippocampal neurons in culture fixed by 100% methanol and labeled for fascin and β-actin. A clear overlap between fascin and β-actin signals can be seen in the growth cone. (**B**) Representative images of neuronal growth cones from cultured DIV2 hippocampal neurons fixed by 4% PFA. While β-actin signals highlight the F-actin bundles in the growth cones, fascin appears to be diffuse and does not highlight F-actin.

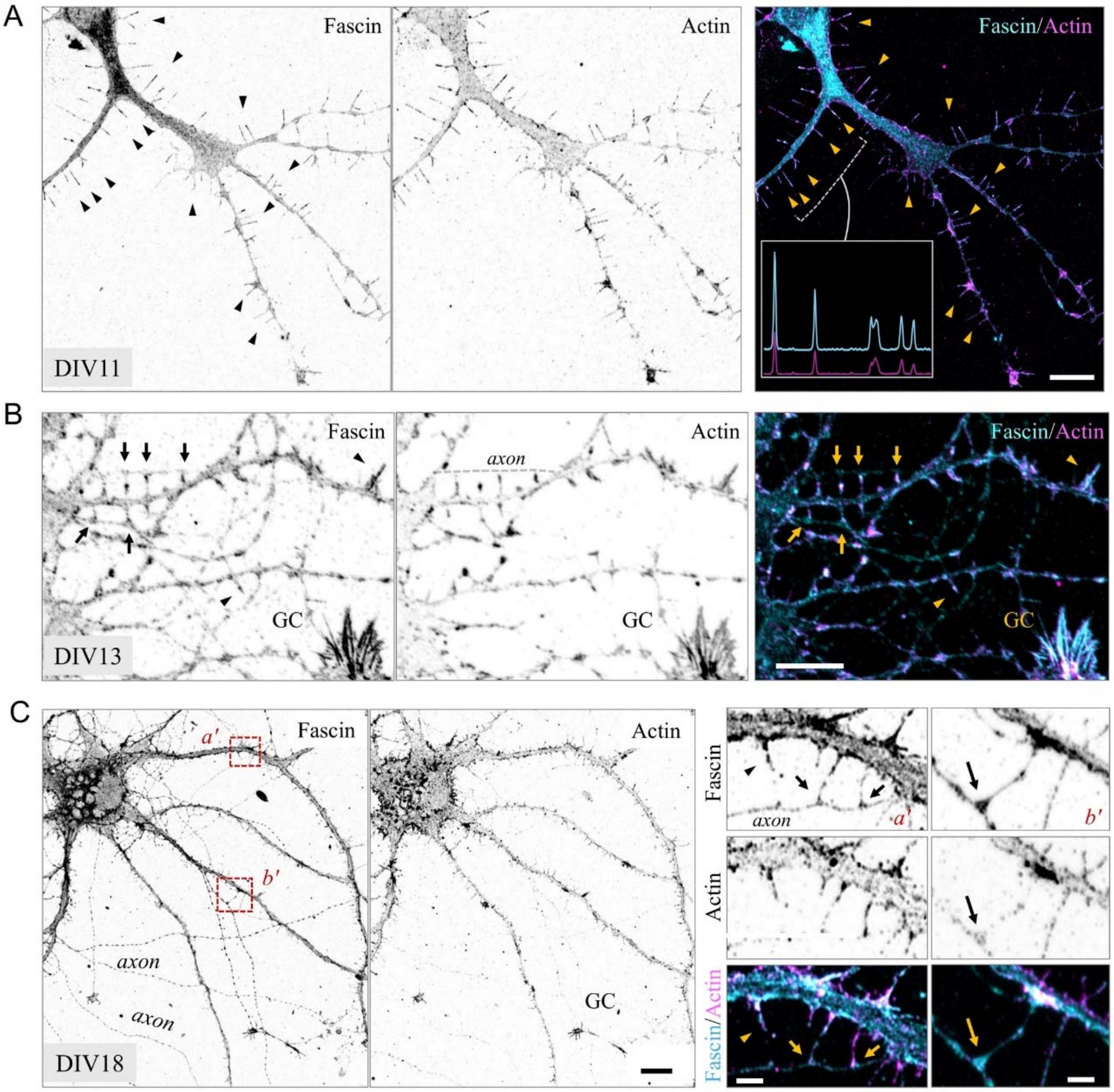


**Figure S4. Fascin is enriched in dendritic filopodia and immature spine-like protrusions during neuronal development.** (**A**) Representative images of low-density cultured hippocampal neurons at DIV11 fixed with 100% cold methanol and labelled for endogenous fascin (cyan) and actin (magenta). The inset in right panel is a linescan showing correlation of fascin and actin levels across multiple filopodia from highlighted region. Arrowheads highlight filopodia with high fascin enrichment. Scale bar = 10 μm. (**B**) Representative images of low-density cultured hippocampal neurons at DIV13 fixed and stained as above. Arrows highlight nascent spines with slightly enlarged heads and fascin enrichment that are in contact with axons. GC: axonal growth cone. Scale bar = 10 μm. (**C**) Representative images of DIV18 neurons fixed and stained as above. Panels on right represent high magnification of corresponding boxed regions from left. Arrows highlight spines with enlarged heads, and arrowheads highlight filopodia. Left scale bar = 10 μm, right scale bars = 2 μm.

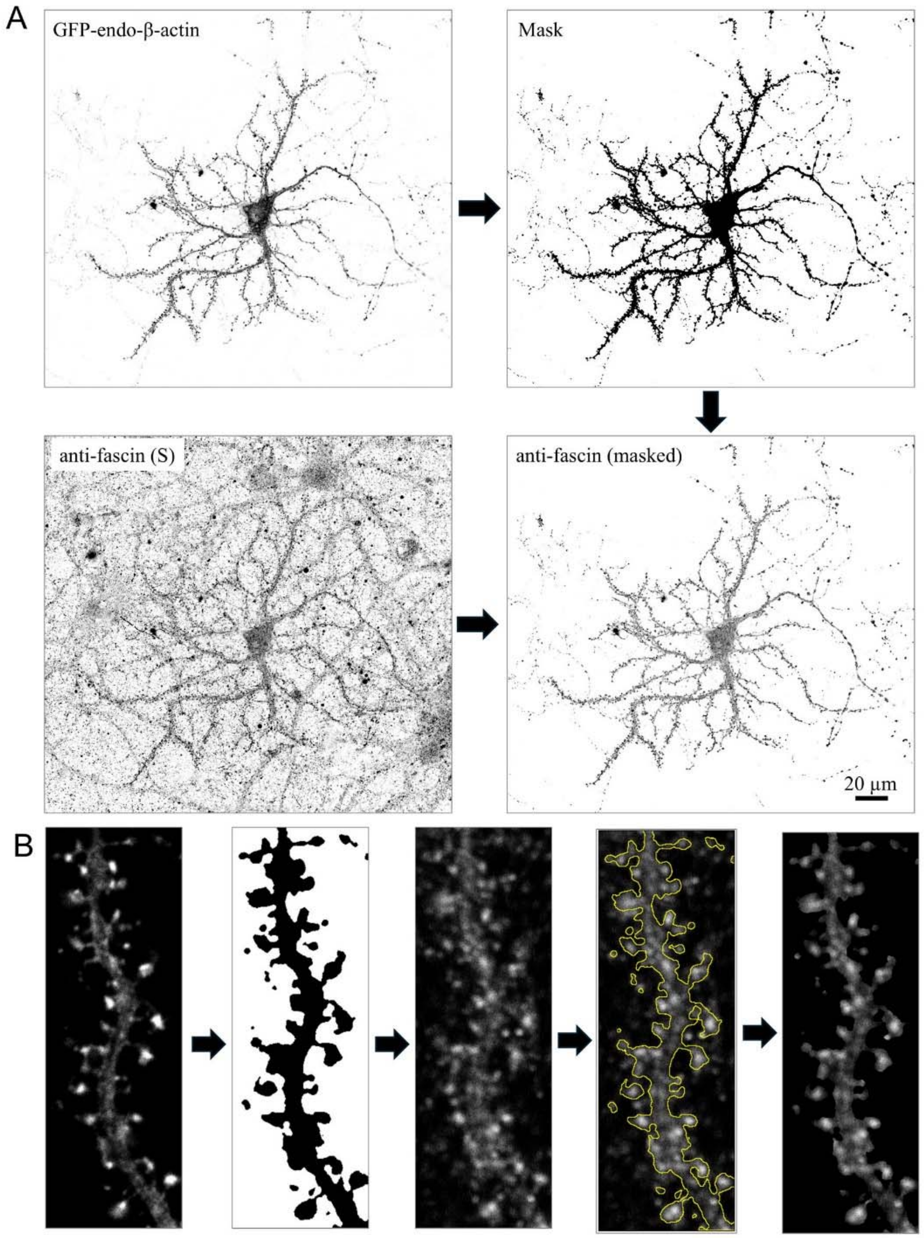


**Figure S5. Image Masking for visualization of dendritic spine localization of fascin in high-density hippocampal cultures.** (**A**) The workflow showing how endogenous GFP-β-actin signal is used to create a binary mask, which is then applied to Sheep anti-fascin staining of high-density hippocampal cultures, in order to isolate fascin signals from individual neurons. (**B**) Representative high-magnification images showing masking and isolation of Sheep anti-fascin staining in dendritic spines.

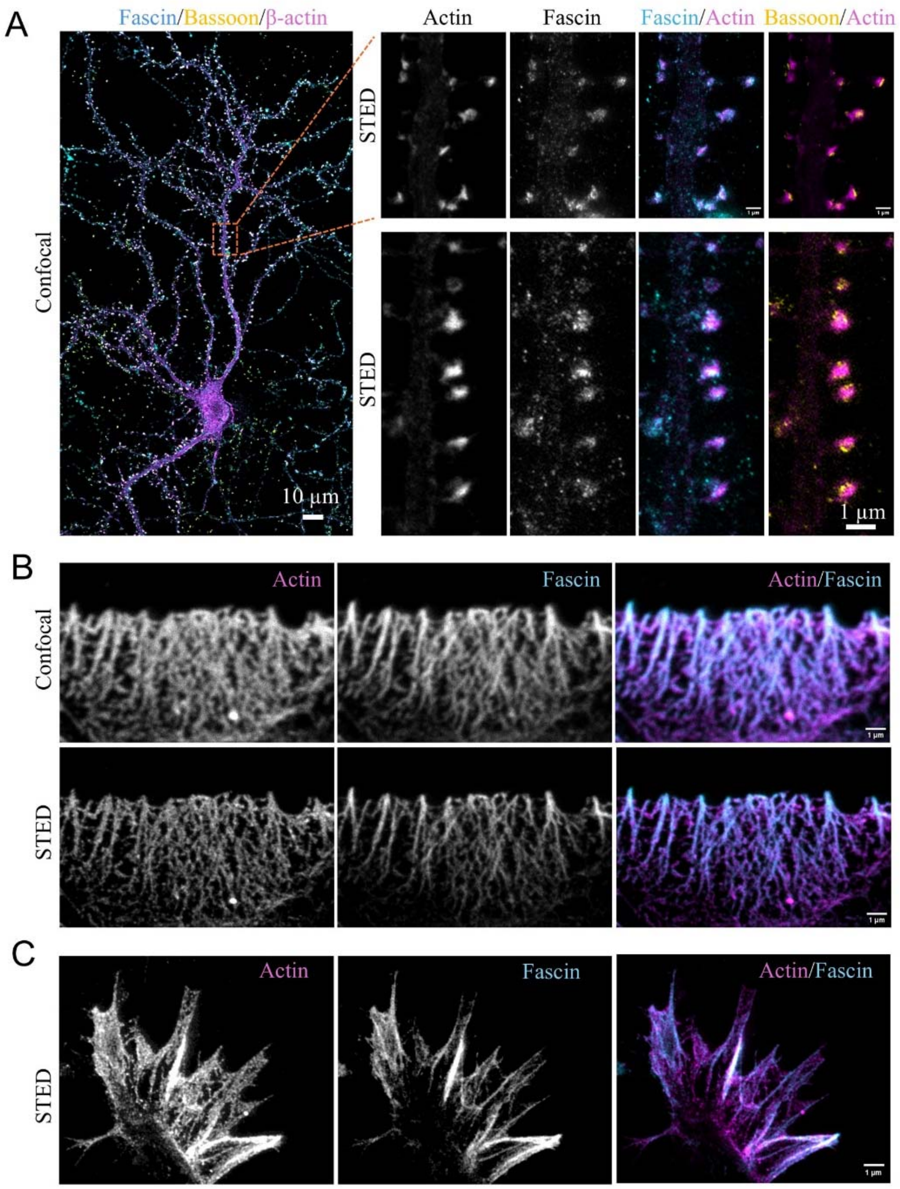


**Figure S6. STED imaging of fascin.** (**A**) Representative confocal image of a DIV25 hippocampal neuron expressing GFP-tagged endogenous β-actin, fixed with 100% cold methanol, and labeled with mouse anti-Bassoon, rabbit anti-GFP, and sheep anti-fascin antibodies. Right, STED images of dendritic regions from the cell on left (top) and an independent cell (bottom), both showing fascin foci in dendritic spine heads. (**B**) Representative confocal and STED images of CAD cell lamellipodia, labelled for endogenous fascin (cyan) and actin (magenta), showing continuous fascin staining pattern along F-actin bundles. (**C**) Representative STED images of growth cones from primary hippocampal neurons, labelled for endogenous fascin (cyan) and actin (magenta), showing continuous fascin staining pattern along F-actin bundles.

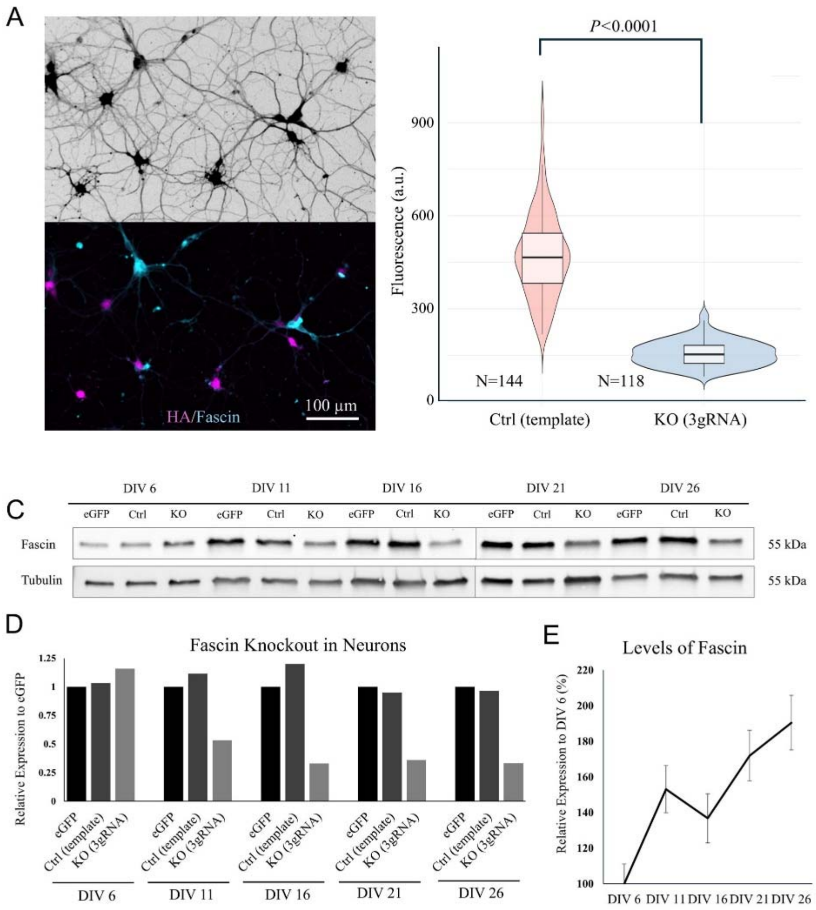


**Figure S7. Fascin1 KO validation in hippocampal neurons.** (**A**) Representative immunofluorescence images of methanol-fixed DIV25 hippocampal neurons that were infected on DIV5 with AAV to express Cas9 and either control (3xgRNA-KO template) or fascin KO (3xgRNA-KO-Fascin1), both of which co-express smFP-HA. Methanol-fixed cells were immunostained for HA to identify transduced cells (magenta) and evaluate fascin levels (cyan). Fascin immunoreactivity is absent in HA-positive fascin KO neurons. (**B**) Violin and box plot showing the loss of fascin fluorescence intensity in KO cells relative to control neurons. N = number of cells analyzed from 3 separate cultures. (**C**) Representative western blots showing the levels of fascin and tubulin (used as a loading control) at various times post AAV infection (i.e. DIV 5). (**D**) Analysis of western blots, showing the effective KO of fascin 12 days post-AAV infection. For each timepoint, samples were normalized to tubulin loading control, and fascin levels in control and KO cells were presented relative to eGFP-expressing neurons. n = 3 separate neuronal cultures. (**E**) Quantification of fascin levels during primary neuron development. Fascin levels were normalized to tubulin and presented relative to the initial DIV6 timepoint. Error bars are ± SEM.
